# The Role of Juvenile Hormone in Midgut Remodeling During *Drosophila melanogaster* Diapause

**DOI:** 10.64898/2026.07.03.736443

**Authors:** Hlib Burtsev, Marc Tatar

## Abstract

Many insects enter diapause, a programmed state of developmental arrest that enables survival under adverse environmental conditions. In *Drosophila melanogaster* Meigen, 1830, diapause is characterized by reproductive arrest and reduced intestinal growth, accompanied by suppressed intestinal stem cell (ISC) activity. Juvenile Hormone (JH) promotes ISC proliferation under favorable conditions, but its capacity to modulate stem cell dynamics during cold-induced diapause remains unclear. Here, we investigated whether JH signaling can reactivate midgut remodeling in adult females maintained at 11℃. At this temperature, flies exhibited pronounced gut atrophy and elevated Phospho-histone H3 (PH3+) cell abundance, consistent with temperature-dependent G2/M phase arrest JH treatment significantly increased the proportion of Delta-positive progenitor cells in the anterior (R2) and posterior (R5) midgut regions at both 11℃ and 25℃, demonstrating that JH acts as a conserved mitogen for the ISC pool irrespective of thermal environment. A trend toward reduced PH3+ accumulation in the posterior midgut following JH treatment (*p* = 0.061) suggests possible facilitation of mitotic exit, though this effect did not reach statistical significance. Despite cellular-level changes, JH treatment did not restore overall gut size, indicating that the 72-84 hour exposure window was insufficient for subsequent tissue hypertrophy. Additionally, we identified a recurrent cold-induced pathology of gut distension, provisionally termed Lumen Obstruction Syndrome (LOS), which was independent of JH signaling. These findings reveal an uncoupling of JH-driven stem cell expansion from gross organ growth under diapause conditions, highlighting the selective sensitivity of the ISC compartment to endocrine signaling during environmental stress.

## INTRODUCTION

### Diapause as an Adaptive Life History Strategy

Many species have evolved adaptations that alter their physiology to survive and reproduce successfully in conditions that are not optimal for them. Diapause is one such strategy and is defined as a genetically programmed state of halted development that animals can enter to postpone their ontogenetic development until favorable conditions return (Koštál, 2006). This study focuses on *Drosophila melanogaster* Meigen, 1830, a cosmopolitan model organism that originated in sub-Saharan Africa, where it adapted to the annual cool, dry seasons of Miombo Woodlands (Pool et al., 2012; Keller, 2007). The genetic plasticity underlying this adaptation enabled many *Drosophila* populations to spread and form locally adapted populations across temperate and tropical regions (Erickson et al., 2020).

Diapause differs from other phenotypically similar life strategies, such as hibernation, estivation, or torpor, in that it causes an organism to alter its physiology and arrest ontogenetic progression. Thus, rather than responding to immediate, often abrupt, environmental cues, diapause ensures an organism’s prolonged lifespan which is sufficient to survive unfavorable conditions (Koštál, 2006). This physiological pivot constitutes a fundamental shift in energy allocation, diverting resources from metabolically expensive vitellogenesis and reproductive maturation toward enhancing somatic maintenance and stress-resistance pathways (Anduaga et al., 2018; Kubrak et al., 2014). Diapausing species are generally divided into two groups of obligate and facultative ones. In obligate organisms, every generation undergoes the diapause period regardless of environmental conditions, as it is a required part of their life history (e.g., the fire bug *Pyrrhocoris apterus,* Linnaeus, 1758; Denlinger, 2002). In contrast, facultative species such as *D. melanogaster* do not need to enter diapause to successfully pass on their genes to the next generation.

Diapause can be conceptually divided into three stages: induction, maintenance, and termination (Koštál, 2006). Induction requires flies not to be fully matured after eclosion (typically, within 6 hours), because if the fly proceeds with the development of ovaries and a fully sized gut, these changes become physiologically irreversible (Kubrak et al., 2014). Therefore, there must be cues in the environment to which the organism responds. In nature, different species and populations of Drosophilidae have evolved sensitivity to photoperiodic and temperature cues in response to their environments (Anduaga et al., 2018). Generally, temperate populations employ a more predictive strategy and are primarily responsive to changes in photoperiod (e.g., *D. montana* and *D. triauraria*), anticipating the upcoming colder season (Lankinen, Kastally and Hoikkala, 2022; Kimura, 1990). In contrast, tropical strains are more responsive to proximate cold stress (Zonato et al., 2017). Beyond light and temperature, recent studies underscore the importance of olfaction in diapause regulation, as specific volatile cues from food, such as yeast, or allelochemicals from plants can act as indicators of seasonal shifts or resource availability, adding a layer of sensory complexity to the induction process (Easwaran & Montell, 2025). In the laboratory, studies usually combine both shortened photoperiod and reduced temperature to maximize diapause induction without artificially imposed diet restriction (Anduaga et al., 2018; Kubrak et al., 2014). During the maintenance phase, the organism enters a state of behavioral and metabolic suppression. Locomotor activity is significantly reduced to conserve energy, yet the fly remains responsive to its environment. Unlike deep torpor, diapausing *Drosophila* must continue to feed to sustain basal metabolic functions, though their nutritional requirements are drastically diminished (Kubrak et al., 2014; Anduaga et al., 2018). This reduced nutrient intake effectively constitutes an endogenous dietary restriction (DR), a condition independently known to extend lifespan at 25°C (Kubrak et al., 2014; Ziehm et al., 2013). As photoperiods lengthen and ambient temperatures return to optimal levels, the endocrine system’s suppression is lifted. These environmental signals trigger a rapid resumption of ontogenetic progression: nutrient uptake increases, and the dormant reproductive and digestive tissues begin their final maturation resulting in the termination of diapause (Saunders et al., 1990; Kubrak et al., 2014; Reiff et al., 2015).

### The Midgut as a Key Tissue in Diapause-Associated Longevity

*D. melanogaster* diapause is primarily studied in the context of adult female arrest of ovarian development and remodeling of the gut. These tissues are among the only major organ systems that contain active stem cells post-eclosion and retain the capacity for significant post-developmental growth, so it is unsurprising that both respond to diapause-inducing conditions by halting their maturation (Easwaran et al., 2022; Singh et al., 2012). Female follicles fail to progress past the last previtellogenic stage (Easwaran et al., 2022). Males are studied less frequently because they lack a readily scorable morphological marker of diapause status analogous to ovarian arrest in females.

In parallel to the reproductive arrest, the largest section of the gut, the midgut, does not undergo the rapid expansion characteristic of normal post-eclosion development. It remains atrophied and shortened until favorable environmental conditions return (Adachi et al., 2025; Reiff et al., 2015). This morphological stasis is maintained by suppressed intestinal stem cell (ISC) proliferation (Adachi et al., 2025). The *Drosophila* midgut is not a homogeneous structure but is partitioned into five distinct functional regions, R1 through R5, each characterized by unique transcriptional profiles and specialized physiological roles (Dutta et al., 2015; Marianes & Spradling, 2013). The posterior regions, R4 and R5, are thought to undergo the most significant remodeling (Adachi et al., 2025). These zones possess a higher density of ISCs and exhibit the greatest sensitivity to nutrient-dependent growth signals, making them the primary drivers of tissue expansion post-diapause (Marianes & Spradling, 2013; Adachi et al., 2025). This quiescent state is characterized by a dramatic reduction in mitotic turnover, quantifiable using the marker Phospho-histone H3 (PH3), and by a diminished pool of actively cycling ISCs, whose identity is marked by expression of the Notch ligand Delta (Dl) (Amcheslavsky et al., 2009; Reiff et al., 2015; Micchelli & Perrimon, 2006). Recent evidence suggests that this mitotic arrest is not a passive consequence of cold stress but is actively regulated by components of the spindle assembly checkpoint (SAC), specifically *BubR1* and *Mad2*. These proteins are proposed to maintain diapausing ISCs in a “primed” G2/M arrested state, preventing the completion of mitosis until favorable conditions return (Adachi et al., 2025).

As flies age, the intestinal epithelium undergoes progressive dysplasia, characterized by a loss of cell polarity and the over-proliferation of ISCs (Jiang & Edgar, 2011; Amcheslavsky et al., 2009). Because the gut is the primary site of both nutrient absorption and the most significant age-related pathology in *Drosophila*, the arrested state of the midgut during diapause is likely a cornerstone of the diapause-associated longevity phenotype (Kubrak et al., 2014; Promislow et al., 1999; Jiang & Edgar, 2011). Studying the gut in this context allows us to observe how a vital organ can postpone its own aging by regulating its internal cellular turnover in response to environmental cues.

### Endocrine Regulation of Diapause: The Role of Juvenile Hormone

A complex interplay between systemic endocrine signals and nutrient-sensing pathways defines the molecular architecture of *D. melanogaster* diapause. The central among these is the Insulin/Insulin-like growth factor signaling (IIS) pathway. Under diapause-inducing conditions, there is a systemic downregulation of *Drosophila* insulin-like peptides (*dILPs*), specifically *dilp2*, *dilp3*, and *dilp5*, which are synthesized in the median neurosecretory cells of the brain (Kubrak et al., 2014). This reduction in IIS activity leads to the upregulation of the forkhead transcription factor FOXO (Tatar et al., 2001; Zečić & Braeckman, 2020). Once active, FOXO translocates to the nucleus to drive the expression of stress-resistance genes, while simultaneously suppressing the Target of Rapamycin (TOR) pathway, further slowing protein synthesis and cellular proliferation (Kubrak et al., 2014).

Juvenile Hormone (JH) is crucial to this endocrine shift (Yamamoto et al., 2013; Saunders et al., 1990). Throughout the life cycle of the fruit fly, JH serves as a critical regulator of developmental timing and tissue maturation (Riddiford et al., 2010; Yamamoto et al., 2013). In larvae, the presence of JH prevents premature metamorphosis, ensuring the organism reaches a critical mass before pupation (Riddiford et al., 2010). In the adult, JH titers rise sharply after eclosion and the hormone transitions to a gonadotropic role, required for the initiation of vitellogenesis in the ovaries and the expansion of the midgut epithelium post-eclosion (Yamamoto et al., 2013; Reiff et al., 2015). In the context of diapause, the corpora allata, the primary site of JH synthesis, enters a quiescent state, resulting in a dramatic drop in systemic JH titers (Yamamoto et al., 2013; Riddiford et al., 2010).

The physiological effects of JH are mediated by its primary receptor, Methoprene-tolerant (Met), a member of the basic helix-loop-helix-Per-Arnt-Sim (bHLH-PAS) family of transcription factors (Charles et al., 2011). Upon binding JH or its analogs, such as methoprene, Met forms a functional complex with the co-activator Taiman (Tai) to regulate the transcription of downstream target genes (Li et al., 2011). In the adult midgut, JH signaling through Met drives gut hypertrophy by stimulating intestinal stem cell (ISC) proliferation via the insulin signaling pathway and the E2F1 transcription factor (Reiff et al., 2015). However, the role of JH in ISC differentiation remains complex. While JH is essential for the rapid expansion of the ISC pool post-mating (Reiff et al., 2015), the subsequent differentiation of progenitors into specialized enterocytes or enteroendocrine cells is governed by local niche signals such as Notch and Delta (Ohlstein & Spradling, 2007). This framework suggests a model in which JH acts as a regulator of stem cell division rate, while other pathways determine final cellular identity (Reiff et al., 2015). This distinction is particularly relevant in the context of the decoupled signaling and mitotic arrest observed in the diapausing midgut (Adachi et al., 2025).

While previous studies have established that the midgut regrows upon return to favorable conditions (Adachi et al., 2025; Reiff et al., 2015), it remains unclear whether this recovery is driven directly by ambient temperature or by the resumption of hormonal signaling. We hypothesize that JH signaling is sufficient to reactivate midgut stem cell dynamics during diapause, even in the continued absence of permissive temperatures. Given that JH stimulates ISC proliferation under normal conditions (Reiff et al., 2015), we predicted that pharmacological application of a JH analog could reactivate dormant stem cells at 11°C. To test this, we maintained flies under constant diapause-inducing conditions (11°C, 8L:16D) while pharmacologically activating JH signaling with methoprene, thereby isolating the hormonal mechanism from environmental factors. This approach allows us to determine whether JH can independently drive midgut remodeling despite the physiological constraints of diapause.

## MATERIALS AND METHODS

### Fly Strains and Husbandry

All experiments were conducted using the wild-type *Drosophila melanogaster* strain white-Dahomey (*wDah*). Stocks were maintained on a standard cornmeal-yeast-agar medium (containing agar, 4% SAF yeast, sugar, cornmeal, and Tegosept). Flies were reared at 25°C under a 12L:12D photoperiod in bottles with the density of parent flies 25 males:25 females per bottle. To ensure the collection of young, virgin females, bottles were moved to 18°C a night prior to eclosion. Eclosing flies were collected within a 6-hr window to ensure their ovaries remained in a pre-vitellogenic state and they could successfully enter diapause.

### Diapause Induction and Hormone Treatment

Experiments were conducted using a 2x2 factorial design to evaluate the independent and interactive effects of temperature (11°C vs. 25°C) and hormone treatment (JH analog (methoprene) vs. EtOH vehicle).

To account for anticipated sample attrition due to the Lumen Obstruction Syndrome (LOS; see Results), two-thirds of the collected virgin females were transferred to an 11°C incubator under a short-day photoperiod (8L:16D) to induce reproductive diapause, while the remaining one-third were maintained at 25°C under a long-day photoperiod (12L:12D). Vials were flipped every two days onto fresh food to maintain optimal nutrition and hygiene. At 22 days (3 weeks) post-induction, half of the flies in each group were treated with either the Juvenile Hormone (JH) analog methoprene (Millipore-Sigma #33375) or an ethanol vehicle control. Treatments were applied by administering 5 µl of solution (0.02 μg methoprene per 1 μl 100% ethanol) onto a fresh synthetic vial plug, following an established protocol (Yamamoto et al., 2013). This method ensured volatile exposure to the hormone while maintaining the flies at 11°C. Flies were exposed to the hormone or vehicle for a duration of 72–84 hours prior to dissection.

Final sample sizes for analysis were: 11℃ JH (n=7), 11℃ EtOH (n=8), 25℃ JH (n=9), 25℃ EtOH (n=12). Flies exhibiting LOS at the time of dissection were excluded from further analysis.

### Immunohistochemistry (IHC)

Midguts were dissected in ice-cold 1x PBS. To ensure tissue integrity, the proventriculus and the midgut-hindgut junction remained intact. Tissues were fixed in 4% paraformaldehyde (PFA) for 20 min at room temperature. Following fixation, samples were washed three times (10 min each) in 0.1% PBST (PBS + 0.1% Triton X-100) and permeabilized in 0.3% PBST for 30 min. Tissues were blocked for 1 hr in a solution of 10% NGS and 2.5% BSA in PBST.

Primary antibody incubation was performed overnight (16–17 hr) at 4°C using rabbit anti-PH3 (1:1000) and mouse anti-Delta (1:50). After three cycles of 10-min washes in PBST, tissues were incubated with secondary antibodies, goat anti-mouse Alexa Fluor 555 (1:200) and goat anti-rabbit Alexa Fluor 647 (1:500), for 1 hr at room temperature. Nuclei were counterstained with DAPI (1:4000, i.e., 0.25 µg/ml) for 10 min. Midguts were mounted in SlowFade Diamond (Invitrogen) on limiting-label slides, sealed with nail polish, and stored in the dark at 4°C until imaging.

### Ovarian Dissection and Histology

Ovaries were collected concurrently with midgut dissections to ensure a matched physiological profile for each biological replicate. Following dissection, ovaries were fixed using the same protocol as the intestinal tissue. For nuclear and actin visualization, samples were incubated for 10 minutes in a combined staining solution of 1:4000 DAPI and 1:400 Phalloidin. Stained ovaries were subsequently mounted in SlowFade Diamond (Invitrogen).

### Imaging, Morphometrics, and Graphics

Midguts were imaged using an Axio Imager.M2 (ZEISS) equipped with a Colibri 7 LED light source. To capture the full extent of the tissue, tiled images were acquired using a 20x objective and processed in ZEN Blue software (ZEISS). Total midgut length and surface area were quantified by defining the central longitudinal axis and the organ perimeter using the spline and contour tools, respectively.

Schematic diagrams were created in GoodNotes (Time Base Technology Ltd.). Figure panels were assembled using Canva (Canva Pty Ltd., Sydney, Australia). Microscopy images were processed in ZEN Blue (ZEISS).

### Cell Quantification and Regional Mapping

For regional analysis, the midgut was partitioned into five functional zones (R1–R5) based on morphological landmarks (Marianes & Spradling, 2013).

*Mitotic Index:* PH3+ cells were quantified manually across the entire volume of the midgut using a 40x objective. All focal planes were scanned to ensure a comprehensive count. For regional comparisons, PH3+ events were categorized as either anterior (R1–R2) or posterior (R4–R5). Notably, mitotic events were consistently absent in the R3 (copper cell) region.

*Stem Cell Density:* Due to non-specific background binding observed with the Delta antibody (DSHB supernatant, clone C594.9B) in our hands, a strict morphology-first quantification pipeline was employed to prevent false positives. First, sampling positions were established in the DAPI channel at the center of each midgut region. Z-stacks were acquired using a 63x objective at these positions and processed into Maximum Intensity Projections (MIPs) (Figure 1). Using the DAPI channel, small nuclei characteristic of stem cells (less than 4–5 µm in diameter) were initially localized (Ohlstein & Spradling, 2006; Micchelli & Perrimon, 2006). Only subsequently was the Delta channel evaluated at those specific coordinates to confirm the presence of characteristic basal punctate staining. To optimize signal-to-noise ratios, brightness and contrast histograms were adjusted locally to distinguish true *Dl*+ puncta from background autofluorescence. Counts were normalized to the total number of DAPI-positive cells in the field of view, which were semi-manually quantified using the point-counting tool in ZEN Blue.

**Figure 1.**
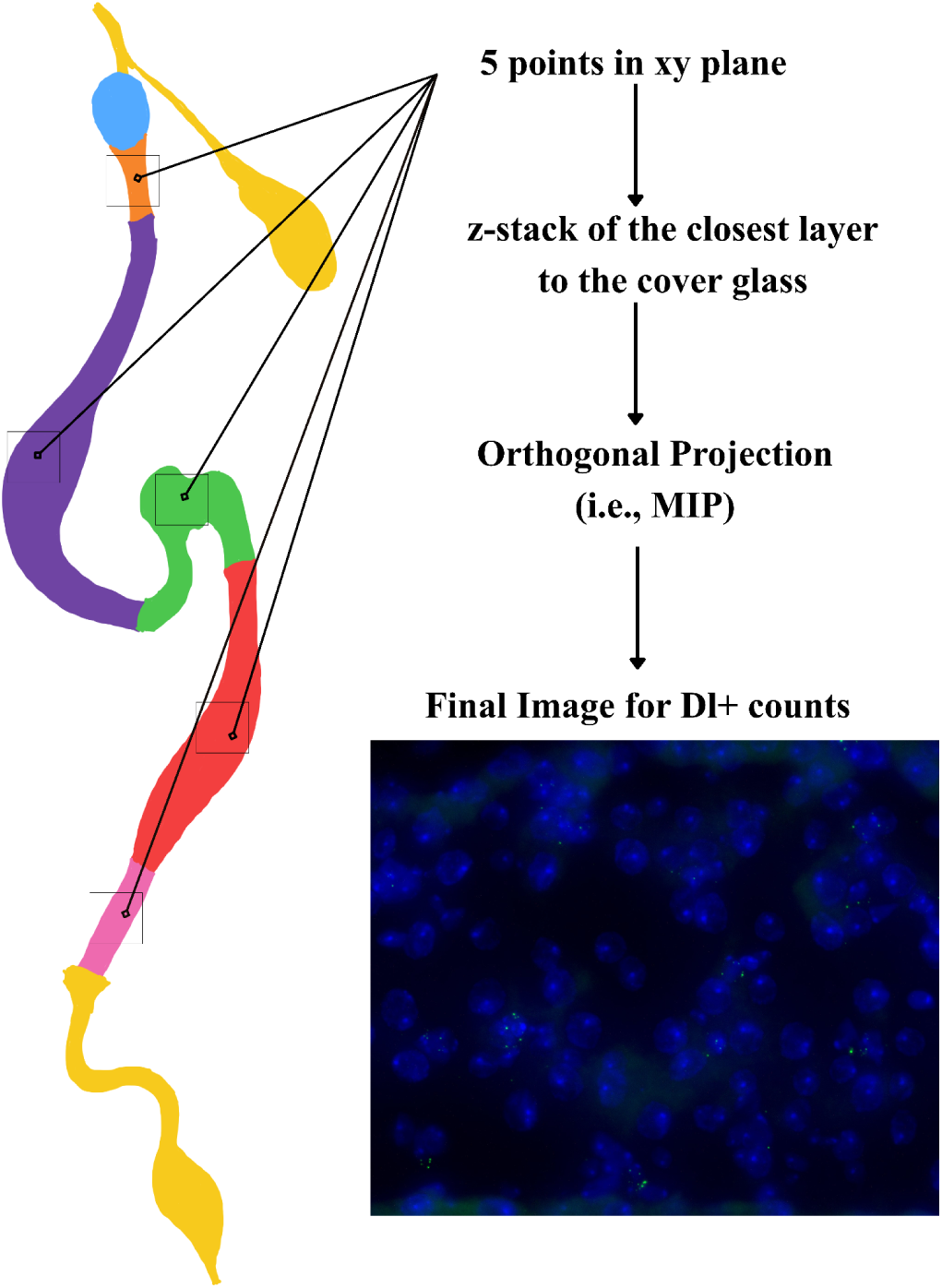
Workflow for imaging Delta+ intestinal stem cells in the midgut across different regions.

### Data Analysis and Statistics

Statistical analyses were performed in R (version 4.5.1) using the tidyverse, rstatix, and betareg packages. For all tests, a *p*-value of < 0.05 was considered statistically significant. Graphical representations of statistical significance in figures are denoted by standard asterisks (* p < 0.05, ** p < 0.01, *** p < 0.001).

For continuous and count data (midgut length, area, and PH3+ cell counts), non-parametric tests were employed given the non-normal distribution of many samples and small sample sizes. To compare all four experimental groups (11C_JH, 11C_EtOH, 25C_JH, 25C_EtOH), a Kruskal-Wallis test was used, followed by Dunn’s post-hoc test with Benjamini-Hochberg (BH) correction for multiple comparisons. Main effects of temperature (11C vs. 25C) and hormone treatment (EtOH vs. JH) were assessed using Wilcoxon rank-sum tests.

Stem cell (*Dl*+) data, expressed as proportions normalized to total DAPI counts, were analyzed using beta regression with a logit link function to account for the bounded nature of the data (0, 1). This model evaluated the main effects of temperature and hormone treatment, as well as their interaction, for each midgut region (R1–R5). Spatial profiles of Dl+ density along the anterior-posterior axis were visualized as mean ± standard error (SEM).

## RESULTS

### Hormonal signaling is sufficient to bypass reproductive arrest at 11°C

To investigate the interplay between environmental temperature and hormonal control in midgut homeostasis, we utilized a *Drosophila melanogaster* model of reproductive diapause (Figure 2). After 3 weeks at 11°C under a short-day photoperiod (8L:16D), females exhibited the expected systemic response to cold-induced arrest, characterized by atrophied, previtellogenic ovaries. To test whether Juvenile Hormone (JH) signaling is sufficient to override this arrest, we treated diapausing flies with a single application of the JH analog methoprene, followed by a 72–84 hour incubation period at 11°C. JH signaling was sufficient to override the temperature-dependent block on oogenesis, resulting in a significant rescue of vitellogenesis despite continued cold exposure (Fisher’s exact test, *p* < 0.001; Figure 2D). This result validated our experimental system and established a baseline for testing whether the midgut, a highly plastic somatic tissue, exhibits comparable sensitivity to JH during diapause.

**Figure 2.**
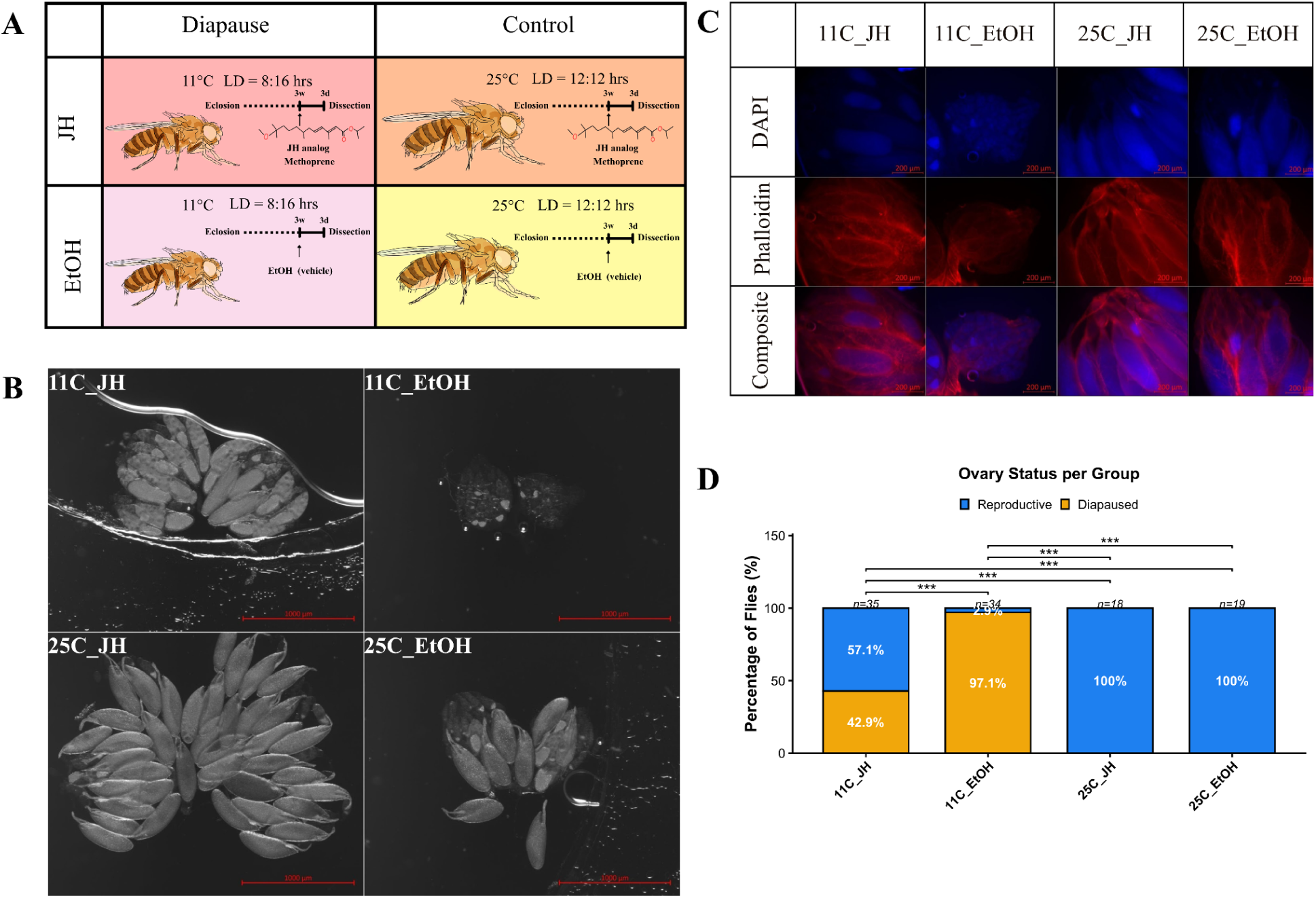
Experimental design, hormone treatment protocol, and ovarian diapause validation. (A) Schematic of the experimental workflow. Newly eclosed wDah females were maintained under diapause-inducing conditions (11°C, 8L:16D) or control conditions (25°C, 12L:12D) for 3 weeks, then treated with either the JH analog methoprene or an ethanol (EtOH) vehicle control for 72–84 hours prior to dissection. (B) Representative brightfield photographs of ovaries from each treatment group. Diapausing females (11°C EtOH) exhibit atrophied, previtellogenic ovaries, whereas JH-treated females (11°C JH) show vitellogenic oocytes comparable to 25°C controls. (C) Widefield fluorescence images of ovaries stained with DAPI (blue, nuclei) and Phalloidin (red, F-actin) across all four treatment groups. (D) Quantification of ovarian developmental status classified as previtellogenic (diapause) or vitellogenic (reproductive). JH treatment significantly rescues vitellogenesis at 11°C (Fisher’s exact test, *p* < 0.001). Sample sizes: 11°C JH (n = 35), 11°C EtOH (n = 34), 25°C JH (n = 18), 25°C EtOH (n = 19).

### Cold temperature induces midgut atrophy and G2/M phase arrest in progenitor cells

After confirming the efficacy of our JH treatment protocol, we examined whether the diapausing midgut exhibits morphological and proliferative changes consistent with developmental arrest. Gross morphometric analysis revealed a significant effect of treatment group on midgut surface area (Kruskal-Wallis, *p* = 0.004; Figure 3C). Post-hoc comparisons confirmed that both 11°C groups exhibited significantly reduced gut area relative to both 25°C groups (Dunn’s post-hoc: 11°C EtOH vs. 25°C JH, *p* = 0.024; 11°C EtOH vs. 25°C EtOH, *p* = 0.021; 11°C JH vs. 25°C JH, *p* = 0.037; 11°C JH vs. 25°C EtOH, *p* = 0.026), while no difference was observed between the two 11°C groups (*p* = 0.914) or between the two 25°C groups (*p* = 0.914). Midgut length followed a similar pattern, with a significant overall effect of temperature (Wilcoxon, *p* = 0.005), though individual pairwise comparisons did not survive correction for multiple testing (Kruskal-Wallis, *p* = 0.046; all Dunn’s post-hoc *p* > 0.05). This atrophy was observed irrespective of hormone treatment (Wilcoxon main effect of drug on area, *p* = 0.987; on length, *p* = 0.671), indicating that the 72–84 hour JH exposure window was insufficient to drive measurable organ-level growth.

To assess the proliferative state of midgut progenitors, we quantified Phospho-histone H3 (PH3)-positive cells, a marker of late G2/M phase, across the entire midgut. The Kruskal-Wallis test revealed a significant effect of the treatment group on total PH3+ counts (*p* = 0.002), as well as within the posterior region (*p* = 0.002) and the anterior region (*p* = 0.045). The main effect of temperature was highly significant: flies maintained at 11°C exhibited substantially elevated PH3+ counts compared to 25°C controls in total (Wilcoxon, *p* = 0.003) and in the posterior midgut (*p* = 0.001). Post-hoc analysis confirmed significant elevations in the 11°C EtOH group relative to both 25°C controls (Dunn’s: 11°C EtOH vs. 25°C JH, total *p* = 0.001; 11°C EtOH vs. 25°C EtOH, total *p* = 0.018).

Rather than indicating heightened proliferation, this accumulation of PH3+ cells at low temperature is consistent with a temperature-dependent deceleration at the G2/M checkpoint, in which cells enter but fail to complete mitosis. PH3+ events were observed predominantly in the anterior (R1–R2) and posterior (R4–R5) regions, with consistent absence from the R3 (copper cell) region.

Within the 11°C cohorts, JH-treated flies exhibited a numerical reduction in PH3+ cell counts relative to ethanol controls. This trend was most pronounced in the posterior midgut (Dunn’s post-hoc, *p* = 0.061), though it did not reach statistical significance at the whole-gut level (*p* = 0.115) or in the anterior region (*p* = 0.825). The main effect of hormone treatment on total PH3+ counts was marginally significant (Wilcoxon, *p* = 0.041), driven primarily by the posterior region, though this collapsed comparison pools both temperature conditions and therefore does not isolate the JH effect within diapause. These data suggest a possible JH-mediated facilitation of mitotic completion in the posterior midgut, but the effect remains inconclusive at the pairwise level given the current sample sizes (11°C JH, n = 7; 11°C EtOH, n = 8).

**Figure 3.**
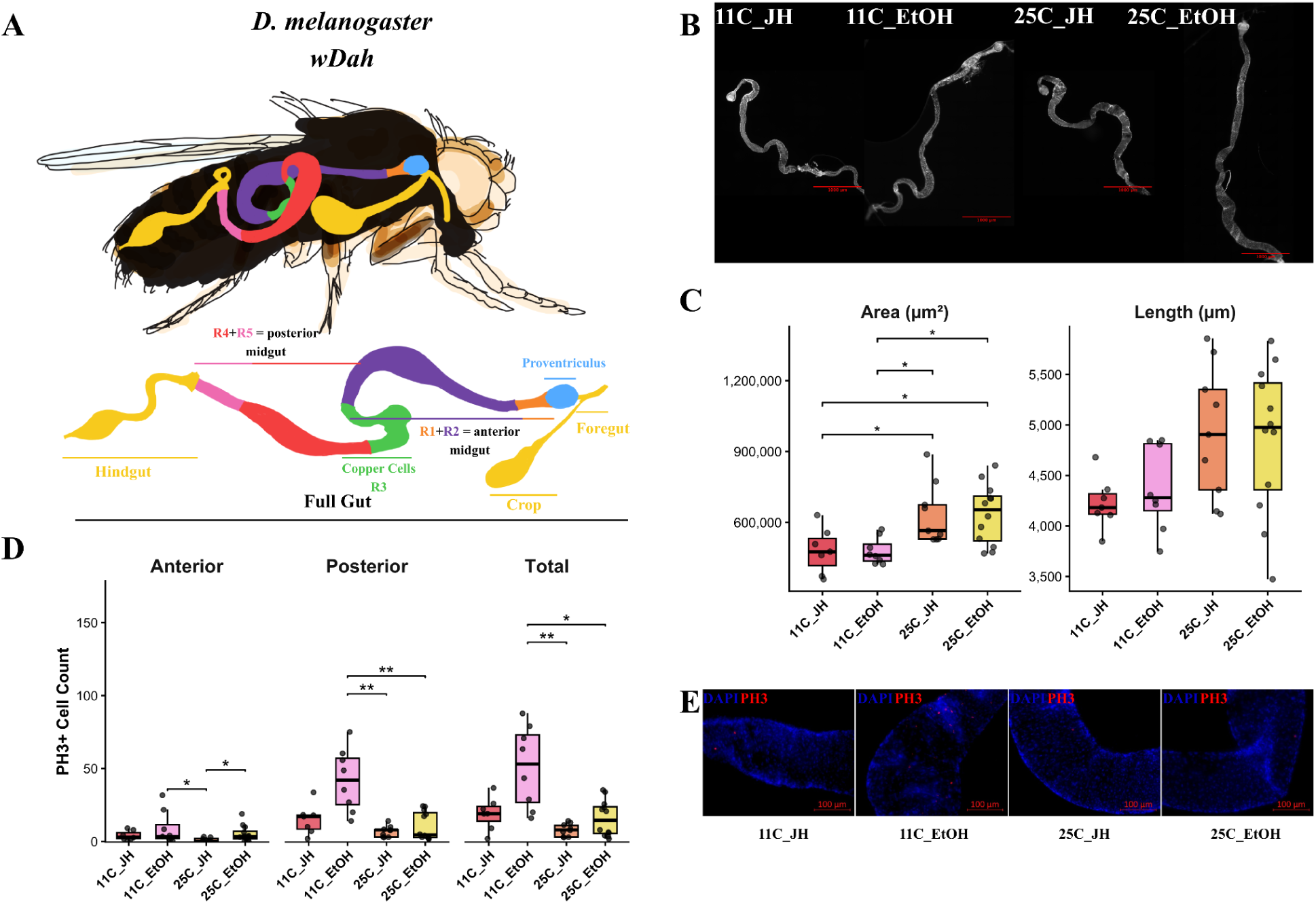
Midgut morphometrics and PH3+ mitotic cell quantification across treatment groups. (A) Anatomical diagram of the *D. melanogaster* midgut illustrating the five functional regions (R1–R5), crop, proventriculus, and hindgut. (B) Representative brightfield photographs of whole midguts from each treatment group, showing the characteristic size reduction associated with 11°C diapause. (C) Quantification of midgut surface area (µm²) and total length (µm). Both 11°C groups exhibit significantly reduced gut area relative to 25°C controls (Kruskal-Wallis, *p* = 0.004; all pairwise 11°C vs. 25°C Dunn’s *p* < 0.05), with no effect of hormone treatment (Wilcoxon main effect of drug, *p* = 0.987). (D) Quantification of PH3+ cells per midgut. Flies at 11°C exhibit significantly elevated PH3+ counts indicative of G2/M phase arrest (Kruskal-Wallis, total *p* = 0.002; posterior *p* = 0.002). Within the 11°C condition, JH treatment produced a numerical reduction in PH3+ counts that approached significance in the posterior midgut (Dunn’s, *p* = 0.061) but was not significant at the whole-gut level (*p* = 0.115). (E) Representative fluorescence images of posterior midgut regions stained for PH3 (red) and DAPI (blue) across all four treatment groups. Sample sizes for cellular analysis: 11°C JH (n = 7), 11°C EtOH (n = 8), 25°C JH (n = 9), 25°C EtOH (n = 12).

### JH signaling induces regional expansion of the intestinal stem cell pool under thermal arrest

Having observed a trend toward reduced PH3+ accumulation following JH treatment, we next asked whether hormonal signaling could expand the progenitor pool itself under diapause conditions. To test this, we quantified the proportion of Delta-positive (Dl+) cells, a marker of intestinal stem cell (ISC) identity, across each of the five midgut regions (R1–R5).

At the whole-gut level (mean Dl+ proportion across all regions), the Kruskal-Wallis test revealed a highly significant effect of the treatment group (*p* < 0.001). The main effect of temperature was significant (Wilcoxon, *p* < 0.001), with 11°C flies maintaining substantially lower Dl+ proportions than 25°C controls. Critically, the main effect of hormone treatment was also significant when collapsed across temperature groups (Wilcoxon, *p* = 0.005), confirming that JH supplementation broadly increased Dl+ proportions regardless of thermal environment. Post-hoc analysis confirmed that the 11°C EtOH group was significantly depleted relative to both 25°C groups (Dunn’s: vs. 25°C JH, *p* < 0.001; vs. 25°C EtOH, *p* = 0.002), while the comparison between 11°C JH and 11°C EtOH trended upward but did not reach significance at the whole-gut level (*p* = 0.087). The 11°C JH group did not differ significantly from the 25°C EtOH group (*p* = 0.209), indicating that JH treatment at 11°C partially rescued the progenitor pool to levels statistically indistinguishable from untreated 25°C controls. However, the 11°C JH group remained significantly lower than the 25°C JH group (*p* = 0.002), indicating that while JH can partially overcome diapause-associated depletion, full rescue requires both hormonal stimulation and a permissive thermal environment.

Regional analysis revealed that the sensitivity to JH is spatially heterogeneous along the anterior–posterior axis. The anterior R2 and posterior R5 regions exhibited the most robust response to JH within the 11°C condition. In R2, JH treatment at 11°C significantly increased the Dl+ proportion relative to 11°C EtOH controls (Dunn’s, *p* = 0.005), effectively rescuing the depletion observed during diapause. The 11°C EtOH group was significantly reduced compared to both 25°C groups (vs. 25°C JH, *p* < 0.001; vs. 25°C EtOH, *p* = 0.009). Similarly, in R5, JH treatment at 11°C significantly expanded the Dl+ pool relative to ethanol controls (Dunn’s, *p* = 0.006), with the 11°C EtOH group again significantly depleted compared to 25°C controls (vs. 25°C JH, *p* < 0.001; vs. 25°C EtOH, *p* = 0.017). Importantly, JH treatment also significantly increased Dl+ proportions at 25°C in both R2 (Dunn’s: 25°C JH vs. 25°C EtOH, *p* = 0.034) and R5 (*p* = 0.006), demonstrating that JH acts as a general mitogen for the ISC pool in these regions irrespective of temperature. This confirms that the JH-mediated expansion observed at 11°C reflects a conserved hormonal mechanism rather than a diapause-specific response, and explains why 11°C JH-treated flies achieve Dl+ levels comparable to 25°C controls: both groups are responding to elevated JH signaling through the same pathway.

Other regions showed more complex or absent responses. In R1, diapause significantly depleted the Dl+ pool (11°C EtOH vs. 25°C JH, *p* = 0.010; vs. 25°C EtOH, *p* < 0.001), but JH treatment at 11°C did not significantly rescue this depletion (*p* = 0.374). In R4, JH treatment at 11°C did not significantly rescue the Dl+ pool relative to ethanol controls (*p* = 0.763), and diapause did not significantly deplete this region relative to 25°C controls (11°C EtOH vs. 25°C EtOH, *p* = 0.086; vs. 25°C JH, *p* = 0.171). However, a highly significant difference was observed between the 25°C JH and 25°C EtOH groups (*p* < 0.001), indicating that R4 is responsive to JH under permissive thermal conditions but not during diapause. The R3 (copper cell) region showed no significant difference between the two 11°C groups, and the overall pattern in this metabolically specialized zone did not conform to the clear depletion-and-rescue pattern observed in R2 and R5. Representative imaging confirmed an expansion of Dl+ clusters in the JH-treated 11°C cohorts compared to ethanol controls, particularly visible in R2 and R5 (Figure 4B). Due to localized mounting or tissue staining artifacts, three regional data points (25C EtOH R1, 25C EtOH R4, 25C EtOH R5) across the entire dataset were omitted from the final analysis (3 out of a total 180 regional observations; see Data Availability Statement).

Together, these results demonstrate that JH acts as a conserved mitogen for the intestinal stem cell pool, capable of driving progenitor expansion at both 25°C and 11°C. However, the capacity for JH to rescue diapause-associated depletion is regionally restricted to R2 and R5 zones that harbor high baseline ISC density under normal conditions and appear preferentially sensitive to endocrine signaling. The significant expansion of the Dl+ pool in R5 parallels the near-significant trend toward reduced PH3+ accumulation in the posterior midgut, providing convergent evidence that the posterior midgut is a primary target of JH action under thermal stress.

**Figure 4.**
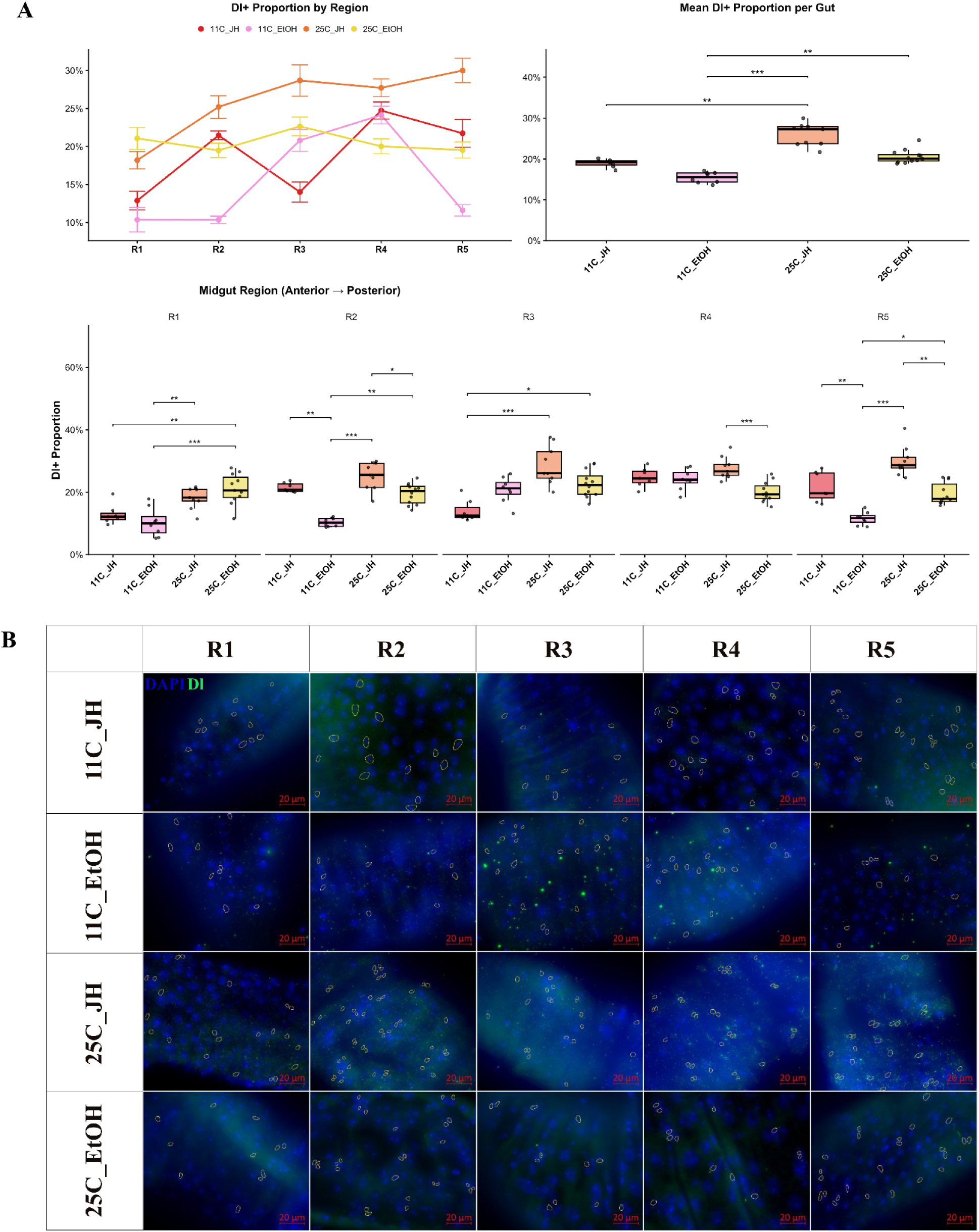
Regional quantification of Delta+ intestinal stem cells along the midgut anterior-posterior axis. (A) Spatial profile of Dl+ cell proportion along the anterior–posterior midgut axis (R1–R5). Mean ± SEM Dl+ proportion is shown per region across treatment groups in the top left. At the whole-gut level, JH treatment significantly increases Dl+ proportion (Wilcoxon main effect, *p* = 0.005), with temperature also significant (Wilcoxon, *p* < 0.001). Regional analysis reveals significant JH-mediated increases within the 11°C condition in R2 (Dunn’s: 11°C JH vs. 11°C EtOH, *p* = 0.005) and R5 (*p* = 0.006). JH also significantly increases Dl+ at 25°C in R2 (*p* = 0.034), R4 (*p* < 0.001) and R5 (*p* = 0.006). (B) Representative widefield fluorescence images of midgut regions R1–R5 stained for Delta (green) and DAPI (blue) across all four treatment groups. Yellow outlines were added during the counting process and indicate individual Dl+ progenitor cells. Note the expanded Dl+ population in 11°C JH and 25°C JH groups relative to their respective EtOH controls, particularly in R2 and R5. Sample sizes: 11°C JH (n = 7), 11°C EtOH (n = 8), 25°C JH (n = 9), 25°C EtOH (n = 12).

### Lumen Obstruction Syndrome (LOS): a diapause-induced mechanical pathology independent of JH signaling

In addition to the cellular changes described above, we identified a recurrent structural pathology during diapause, which we provisionally termed Lumen Obstruction Syndrome (LOS). This phenotype was characterized by severe midgut distension localized to the R2 region, attributable to the retention of meconium and subsequent accumulation of luminal contents (Figure 5). In severe cases, the distended gut wall ruptured during dissection, revealing compacted material within the lumen (Figure 5A). Representative images and schematic diagrams of progressively distended midguts are shown in Figure 5A-C.

The incidence of LOS was dramatically elevated at 11°C. Fisher’s exact tests confirmed significant differences between 11°C groups and their 25°C counterparts: 11°C EtOH vs. 25°C EtOH (p < 0.001), 11°C EtOH vs. 25°C JH (p = 0.042), 11°C JH vs. 25°C JH (p < 0.001), and 11°C JH vs. 25°C EtOH (p < 0.001). Specifically, 62.9% of the 11°C JH-treated cohort and 44.1% of the 11°C ethanol controls exhibited severe LOS, compared to only 11.1% of the 25°C JH-treated flies and 0% of the 25°C ethanol controls (Figure 5D).

JH treatment did not mitigate LOS. Rather, it trended toward exacerbating the phenotype at 11°C (62.9% vs. 44.1%), although this difference between JH and EtOH within the 11°C group was not significant (Fisher’s exact, p = 0.180). Logistic regression restricted to the 11°C cohorts confirmed no significant independent effect of hormone treatment on LOS incidence (*p* = 0.242; Figure 5D). Flies exhibiting severe LOS were excluded from all cellular quantification (PH3 and Dl+ analyses) to prevent confounding effects of mechanical disruption on immunohistochemistry.

These data indicate that while JH can reactivate the cellular machinery of the midgut, it is insufficient to resolve the gross mechanical consequences of cold exposure. The trend toward elevated LOS incidence in JH-treated flies may reflect increased metabolic demand or feeding behavior in a digestive tract that remains functionally obstructed, though this interpretation remains speculative given the non-significant difference.

**Figure 5.**
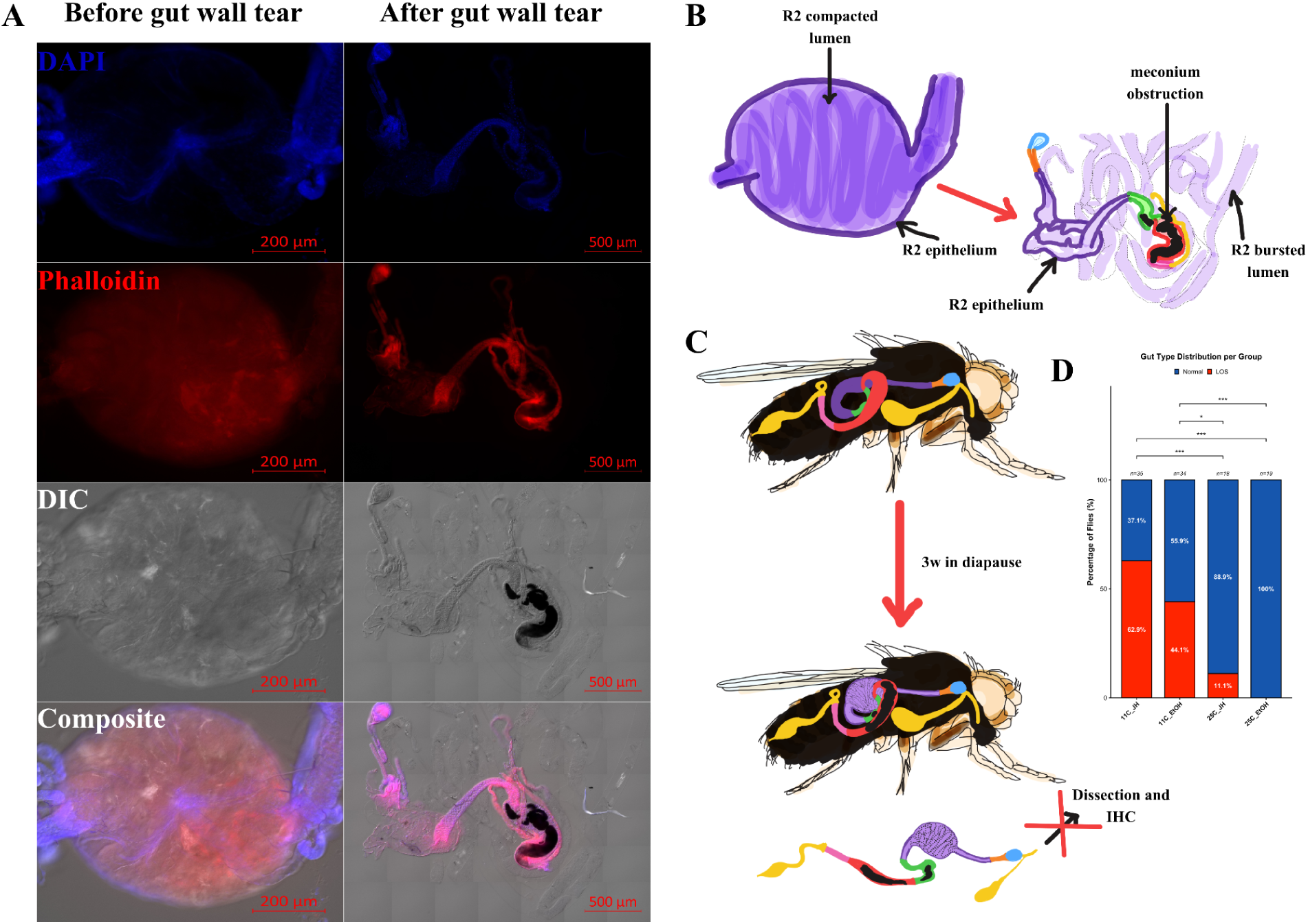
Characterization of Lumen Obstruction Syndrome (LOS) during diapause. (A) Fluorescence images of the R2 midgut region before and after gut wall rupture during dissection, stained with DAPI (blue), Phalloidin (red), and imaged under DIC. Compacted luminal contents are visible following epithelial breach. (B) Schematic illustration of the structures from photographs in subfigure (A). (C) Anatomical schematic illustrating the structural basis of LOS: meconium retained post-eclosion obstructs the lumen, with subsequent food accumulation causing progressive distension of the R2 epithelium over 3 weeks of diapause. (D) Frequency distribution of gut phenotypes (Normal vs. LOS) across treatment groups. LOS incidence is significantly elevated at 11°C (Fisher’s exact: 11°C EtOH vs. 25°C EtOH, *p* < 0.001; 11°C JH vs. 25°C JH, *p* < 0.001). JH did not significantly affect LOS incidence within the 11°C condition (Fisher’s exact, *p* = 0.180). Percentages: 11°C JH = 62.9% LOS, 11°C EtOH = 44.1%, 25°C JH = 11.1%, 25°C EtOH = 0%. Sample sizes represent total flies dissected prior to exclusion criteria and are 11°C JH (n = 35), 11°C EtOH (n = 34), 25°C JH (n = 18), 25°C EtOH (n = 19).

## DISCUSSION

In this study, we investigated whether Juvenile Hormone (JH) signaling can reactivate midgut remodeling in *Drosophila melanogaster* during cold-induced reproductive diapause. By maintaining flies under constant diapause-inducing conditions (11°C, 8L:16D) while pharmacologically activating JH signaling, we isolated the hormonal mechanism from environmental factors that normally accompany diapause termination. Our results demonstrate that JH acts as a conserved mitogen for the intestinal stem cell pool that is capable of driving regional expansion of Delta-positive progenitors at both permissive and restrictive temperatures. However, this cellular reactivation was not accompanied by gross organ-level growth, revealing a potential fundamental uncoupling between stem cell proliferation and tissue hypertrophy under thermal stress.

### JH as a conserved intestinal stem cell mitogen

The main finding of this study is that JH signaling significantly expanded the Dl+ progenitor pool in the anterior R2 and posterior R5 midgut regions at 11°C, effectively rescuing the diapause-associated depletion to levels statistically indistinguishable from untreated 25°C controls. Critically, JH also drove significant Dl+ expansion at 25°C in the same regions, demonstrating that this is a conserved hormonal mechanism rather than a diapause-specific response. This finding is consistent with the established role of JH signaling through Met in stimulating ISC proliferation via the insulin signaling pathway and the E2F1 transcription factor (Reiff et al., 2015), and extends this framework by demonstrating that the pathway remains functional even under the severe metabolic constraints of diapause.

The regional specificity of this response is quite notable. R2 and R5 are zones known to harbor high baseline ISC density (Marianes & Spradling, 2013) and R4-R5 were shown to undergo the most significant remodeling during diapause recovery (Adachi et al., 2025). Our data suggest that these regions possess a “privileged” sensitivity to systemic endocrine signals, potentially due to higher expression of Met or downstream pathway components. In contrast, R1 showed significant depletion during diapause but was refractory to JH rescue, suggesting that the molecular competence to respond to JH is not uniform along the anterior-posterior axis. The R3 copper cell region showed no clear diapause-associated depletion or JH response, consistent with its specialized metabolic identity and distinct regulatory programs (Dutta et al., 2015; Marianes & Spradling, 2013).

Interestingly, R4 exhibited a significant JH response at 25°C (25°C JH vs. 25°C EtOH, *p* < 0.001) but not at 11°C (*p* = 0.763), suggesting that this region is competent to respond to JH under permissive thermal conditions but that cold temperature specifically abolishes this responsiveness. This contrasts with R2 and R5, where JH drove significant expansion at both temperatures. The temperature-dependent loss of JH sensitivity in R4 may reflect region-specific differences in the thermodynamic requirements for ISC activation, or a higher threshold of JH signaling needed to overcome diapause-associated suppression in this zone. Future studies should investigate whether R4 ISCs express lower levels of Met or downstream pathway components at 11°C compared to R2 and R5.

### Uncoupling of stem cell proliferation from organ-level growth

Another central finding of this study is the apparent paradox between the cellular and morphological responses to JH at 11°C. While JH supplementation drove significant expansion of the Dl+ progenitor pool in R2 and R5 and produced a near-significant trend toward reduced PH3+ accumulation in the posterior midgut (*p* = 0.061), there was no significant change in overall midgut length or surface area between JH-treated and control flies at either temperature.

Several factors likely contribute to this dissociation. First, the temporal window of hormone exposure (72–84 hours) may be sufficient to trigger transcriptional changes and initiate progenitor expansion, but insufficient for the subsequent steps of cellular differentiation into mature enterocytes and physical hypertrophy of the gut epithelium. Enterocyte differentiation from ISC progeny requires Notch-mediated fate decisions (Ohlstein & Spradling, 2007) and subsequent endoreplication to achieve the large cell volumes characteristic of mature absorptive cells, processes that likely require more than three days to complete. This temporal limitation is supported by the absence of significant morphological differences between the 25°C JH and 25°C EtOH groups, suggesting that even under optimal thermal conditions, 72–84 hours is insufficient for JH to drive gross tissue remodeling.

Second, the 11°C environment likely imposes severe thermodynamic constraints on cellular growth independently of hormonal signaling. While JH may provide the transcriptional signal to divide, the biophysical processes of protein synthesis, membrane expansion, and cytoskeletal rearrangement that are required for cell growth operate at drastically reduced rates at 11°C. This interpretation is consistent with the framework proposed by Adachi et al. (2025), in which BubR1 and Mad2 maintain ISCs in a primed G2/M state during diapause. Our data suggest that JH signaling may partially override this checkpoint in the posterior midgut, as indicated by the near-significant reduction in PH3+ cells (*p* = 0.061), but the completion of subsequent growth steps remains thermally constrained.

Together, these results suggest that JH specifically targets ISC identity and proliferative capacity, acting as a direct mitogen for the stem cell compartment, whereas downstream organ-level growth requires either extended temporal windows or a permissive thermal environment to proceed.

### PH3 accumulation as evidence of temperature-dependent G2/M deceleration

The elevated PH3+ counts observed at 11°C represent one of the clearest morphological signatures of diapause in the midgut. Rather than indicating heightened proliferative activity, this accumulation is consistent with a model in which progenitor cells enter the G2/M phase but cannot complete mitosis at low temperature. This interpretation aligns with the recently proposed role of spindle assembly checkpoint components BubR1 and Mad2 in actively maintaining diapausing ISCs in a G2/M arrested state (Adachi et al., 2025).

The trend toward reduced PH3+ counts following JH treatment at 11°C (*p* = 0.061 in the posterior midgut) is suggestive of JH-mediated facilitation of mitotic exit, but we emphasize that this effect did not reach significance at the pairwise level. The marginally significant main effect of hormone treatment on total PH3+ counts (Wilcoxon, *p* = 0.041) must be interpreted cautiously, as this collapsed comparison pools both temperature groups and therefore does not isolate the JH effect within the diapause condition specifically. Larger sample sizes will be required to definitively determine whether JH can override the SAC-mediated G2/M arrest or whether the observed trend reflects stochastic variation in a small cohort.

Nevertheless, the convergence of two independent observations in the posterior midgut, namely the significant expansion of the Dl+ pool in R5 (*p* = 0.006) alongside the near-significant reduction in PH3+ cells (*p* = 0.061), provides convergent evidence that this region is preferentially sensitive to JH signaling under thermal stress.

### Lumen Obstruction Syndrome: a novel diapause-induced mechanical pathology

In addition to the cellular effects of JH on the stem cell compartment, we identified a recurrent structural pathology during diapause, provisionally termed Lumen Obstruction Syndrome (LOS). The high incidence of LOS at 11°C (44–63% of flies) represents a significant source of sample attrition and a previously undescribed consequence of prolonged cold exposure in laboratory diapause assays.

We hypothesize that LOS results from an initial failure to excrete larval meconium post-eclosion, likely due to the immediate transfer of newly eclosed flies to 11°C before the midgut has achieved sufficient peristaltic competence. Because diapausing flies continue to feed at reduced rates to sustain basal metabolism (Kubrak et al., 2014), newly ingested material progressively accumulates behind this initial obstruction over the three-week diapause period. The resulting mechanical distension of the R2 epithelium beyond its elastic limits explains the gut wall rupture observed during dissection.

Importantly, JH treatment did not prevent or mitigate LOS (Fisher’s exact, *p* = 0.180 between 11°C groups), confirming that this pathology operates independently of the hormonal signaling axis investigated here. The trend toward higher LOS incidence in JH-treated flies (62.9% vs. 44.1%), while not significant, raises the possibility that JH-mediated increases in metabolic demand or feeding behavior could accelerate luminal accumulation in a functionally obstructed tract, though this remains speculative.

LOS has important methodological implications for future diapause studies. Researchers employing prolonged cold exposure protocols should anticipate substantial sample attrition and design experiments accordingly if they work with *wDah* background and, potentially, others. The exclusion of LOS-affected guts from our cellular analyses introduces a potential survivorship bias: the flies we analyzed represent the subset that successfully cleared their meconium or avoided obstruction, and may therefore not fully represent the whole diapausing population.

Future studies employing dye-tracking assays, histological cross-sections, or time-course analyses of LOS onset will be valuable for characterizing the temporal dynamics and compositional nature of this obstruction.

### Technical limitations and future directions

Several methodological limitations should be acknowledged when interpreting these results. First, the sample sizes (n = 7–12 per group) limit statistical power, particularly for the PH3 analyses where the JH effect within 11°C did not reach significance. The unequal group sizes reflect the anticipated attrition from LOS, and while we planned for this with our 2:1 allocation ratio, the final analyzed cohorts remain small by conventional standards.

Second, the quantification of the ISC pool was technically challenging due to the non-specific background binding observed with the Delta antibody (DSHB supernatant, clone C594.9B) in our preparations. Our morphology-first quantification pipeline, which required initial identification of small nuclei in the DAPI channel before evaluating the Delta signal, represents a conservative approach that likely underestimates the true Dl+ population. Despite this conservative bias, we still detected significant regional effects, suggesting that the true biological signal may be even stronger than reported.

Third, we cannot rule out the possibility that methoprene exposure at 11°C triggers secondary signaling cascades beyond direct Met-Tai transcriptional activation. JH is known to interact with the IIS pathway (Reiff et al., 2015; Yamamoto et al., 2013), and the observed ISC expansion could reflect indirect effects through insulin signaling rather than direct JH-Met action in the midgut. Genetic approaches using tissue-specific Met knockdown would be necessary to distinguish between these possibilities.

Future studies should extend the hormone exposure window beyond 72–84 hours to determine whether prolonged JH signaling at 11°C can eventually drive measurable tissue hypertrophy, or whether cold temperature represents an absolute constraint on organ growth regardless of hormonal status. Additionally, combining JH treatment with a brief temperature shift (e.g., 11°C to 25°C for 24 hours following JH exposure) could help dissect the relative contributions of hormonal priming versus thermal permissiveness to the completion of tissue remodeling.

## CONCLUSION

This study demonstrates that JH signaling retains the capacity to reactivate intestinal stem cell dynamics during cold-induced diapause in *D. melanogaster*. By isolating the hormonal mechanism from the environmental cues that normally accompany diapause termination, we show that the stem cell compartment remains acutely sensitive to systemic JH even at 11°C. However, the translation of stem cell expansion into organ-level growth remains constrained by low temperature, revealing a hierarchical model in which JH acts as the initial endocrine trigger for midgut remodeling, while ambient temperature gates the downstream biophysical processes required for tissue hypertrophy. This uncoupling of proliferation from growth highlights the stem cell niche as a selectively sensitive node within the broader diapause regulatory network, and identifies R2 and R5 as primary regional targets of JH action in the adult midgut.

## ACKNOWLEDGEMENTS

We are deeply grateful to the Brown Center for the Biology of Aging for their vital institutional support. We sincerely thank the members of the Silva García Lab for their immense generosity in providing access to their specialized imaging facilities and for their invaluable technical expertise. The high-resolution microscopy in this study would not have been possible without their collaborative spirit. Finally, we thank our colleagues and lab members for their hands-on assistance during dissections and for the critical discussions that greatly improved this manuscript.

## DATA AVAILABILITY STATEMENT

The data and code that support the findings of this study are openly available in Zenodo at https://doi.org/10.5281/zenodo.21094697. A comprehensive summary table detailing all statistical comparisons and analyses is provided in the Supporting Information.

## AUTHOR CONTRIBUTIONS

HB: Conceptualization, Methodology, Formal Analysis, Investigation, Writing, Visualization. MT: Conceptualization, Resources, Supervision, Review.

