## Supporting Information for "The Role of Juvenile Hormone in Midgut Remodeling During *Drosophila melanogaster* Diapause"

### PH3+ Count

| Test | Subgroup | Comparison | Statistic | p (BH−adj) | Sig |
|---|---|---|---|---|---|
| Dunn (BH) | Anterior | 11C_EtOH vs 25C_EtOH | −0.294 | 8.25e−01 | ns |
| Dunn (BH) | Anterior | 11C_EtOH vs 25C_JH | −2.464 | 4.77e−02 | * |
| Dunn (BH) | Anterior | 11C_JH vs 11C_EtOH | 0.463 | 8.25e−01 | ns |
| Dunn (BH) | Anterior | 11C_JH vs 25C_EtOH | 0.221 | 8.25e−01 | ns |
| Dunn (BH) | Anterior | 11C_JH vs 25C_JH | −1.901 | 1.15e−01 | ns |
| Dunn (BH) | Anterior | 25C_JH vs 25C_EtOH | 2.411 | 4.77e−02 | * |
| Dunn (BH) | Posterior | 11C_EtOH vs 25C_EtOH | −3.384 | 2.15e−03 | ** |
| Dunn (BH) | Posterior | 11C_EtOH vs 25C_JH | −3.518 | 2.15e−03 | ** |
| Dunn (BH) | Posterior | 11C_JH vs 11C_EtOH | 2.162 | 6.12e−02 | ns |
| Dunn (BH) | Posterior | 11C_JH vs 25C_EtOH | −0.894 | 4.45e−01 | ns |
| Dunn (BH) | Posterior | 11C_JH vs 25C_JH | −1.172 | 3.62e−01 | ns |
| Dunn (BH) | Posterior | 25C_JH vs 25C_EtOH | 0.375 | 7.08e−01 | ns |
| Dunn (BH) | Total | 11C_EtOH vs 25C_EtOH | −2.741 | 1.84e−02 | * |
| Dunn (BH) | Total | 11C_EtOH vs 25C_JH | −3.696 | 1.31e−03 | ** |
| Dunn (BH) | Total | 11C_JH vs 11C_EtOH | 1.898 | 1.15e−01 | ns |
| Dunn (BH) | Total | 11C_JH vs 25C_EtOH | −0.565 | 5.72e−01 | ns |
| Dunn (BH) | Total | 11C_JH vs 25C_JH | −1.615 | 1.60e−01 | ns |
| Dunn (BH) | Total | 25C_JH vs 25C_EtOH | 1.236 | 2.60e−01 | ns |
| Kruskal−Wallis | Anterior | All 4 groups | 8.029 | 4.54e−02 | * |
| Kruskal−Wallis | Posterior | All 4 groups | 15.268 | 1.60e−03 | ** |
| Kruskal−Wallis | Total | All 4 groups | 14.346 | 2.47e−03 | ** |
| Wilcoxon rank−sum | Anterior | 11C vs 25C | 201 | 1.62e−01 | ns |
| Wilcoxon rank−sum | Anterior | EtOH vs JH | 224.5 | 3.90e−02 | * |
| Wilcoxon rank−sum | Posterior | 11C vs 25C | 258 | 1.30e−03 | ** |
| Wilcoxon rank−sum | Posterior | EtOH vs JH | 209 | 1.22e−01 | ns |
| Wilcoxon rank−sum | Total | 11C vs 25C | 252 | 2.53e−03 | ** |
| Wilcoxon rank−sum | Total | EtOH vs JH | 224.5 | 4.14e−02 | * |

### Morphology

| Test | Subgroup | Comparison | Statistic | p (BH−adj) | Sig |
|---|---|---|---|---|---|
| Dunn (BH) | Area (µm²) | 11C_EtOH vs 25C_EtOH | 2.92 | 2.10e−02 | * |
| Dunn (BH) | Area (µm²) | 11C_EtOH vs 25C_JH | 2.645 | 2.45e−02 | * |
| Dunn (BH) | Area (µm²) | 11C_JH vs 11C_EtOH | −0.291 | 9.14e−01 | ns |
| Dunn (BH) | Area (µm²) | 11C_JH vs 25C_EtOH | 2.485 | 2.59e−02 | * |
| Dunn (BH) | Area (µm²) | 11C_JH vs 25C_JH | 2.251 | 3.66e−02 | * |
| Dunn (BH) | Area (µm²) | 25C_JH vs 25C_EtOH | 0.108 | 9.14e−01 | ns |
| Dunn (BH) | Length (µm) | 11C_EtOH vs 25C_EtOH | 1.759 | 1.43e−01 | ns |
| Dunn (BH) | Length (µm) | 11C_EtOH vs 25C_JH | 1.668 | 1.43e−01 | ns |
| Dunn (BH) | Length (µm) | 11C_JH vs 11C_EtOH | 0.547 | 7.01e−01 | ns |
| Dunn (BH) | Length (µm) | 11C_JH vs 25C_EtOH | 2.283 | 8.99e−02 | ns |
| Dunn (BH) | Length (µm) | 11C_JH vs 25C_JH | 2.17 | 8.99e−02 | ns |
| Dunn (BH) | Length (µm) | 25C_JH vs 25C_EtOH | −0.018 | 9.86e−01 | ns |
| Kruskal−Wallis | Area (µm²) | All 4 groups | 13.595 | 3.51e−03 | ** |
| Kruskal−Wallis | Length (µm) | All 4 groups | 8.003 | 4.59e−02 | * |
| Wilcoxon rank−sum | Area (µm²) | 11C vs 25C | 43 | 1.06e−04 | *** |
| Wilcoxon rank−sum | Area (µm²) | EtOH vs JH | 161 | 9.87e−01 | ns |
| Wilcoxon rank−sum | Length (µm) | 11C vs 25C | 71 | 4.75e−03 | ** |
| Wilcoxon rank−sum | Length (µm) | EtOH vs JH | 174 | 6.71e−01 | ns |

### DI+ Proportion

| Test | Subgroup | Comparison | Statistic | p (BH−adj) | Sig |
|---|---|---|---|---|---|
| Dunn (BH) | R1 | 11C_EtOH vs 25C_EtOH | 3.914 | 5.44e−04 | *** |
| Dunn (BH) | R1 | 11C_EtOH vs 25C_JH | 2.89 | 9.90e−03 | ** |
| Dunn (BH) | R1 | 11C_JH vs 11C_EtOH | −0.889 | 3.74e−01 | ns |
| Dunn (BH) | R1 | 11C_JH vs 25C_EtOH | 2.81 | 9.90e−03 | ** |
| Dunn (BH) | R1 | 11C_JH vs 25C_JH | 1.874 | 9.15e−02 | ns |
| Dunn (BH) | R1 | 25C_JH vs 25C_EtOH | 0.922 | 3.74e−01 | ns |
| Dunn (BH) | R2 | 11C_EtOH vs 25C_EtOH | 2.851 | 8.72e−03 | ** |
| Dunn (BH) | R2 | 11C_EtOH vs 25C_JH | 4.743 | 1.27e−05 | *** |
| Dunn (BH) | R2 | 11C_JH vs 11C_EtOH | −3.17 | 4.57e−03 | ** |
| Dunn (BH) | R2 | 11C_JH vs 25C_EtOH | −0.714 | 4.75e−01 | ns |
| Dunn (BH) | R2 | 11C_JH vs 25C_JH | 1.317 | 2.25e−01 | ns |
| Dunn (BH) | R2 | 25C_JH vs 25C_EtOH | −2.275 | 3.43e−02 | * |
| Dunn (BH) | R3 | 11C_EtOH vs 25C_EtOH | 0.507 | 6.12e−01 | ns |
| Dunn (BH) | R3 | 11C_EtOH vs 25C_JH | 2.234 | 5.09e−02 | ns |
| Dunn (BH) | R3 | 11C_JH vs 11C_EtOH | 2.108 | 5.26e−02 | ns |
| Dunn (BH) | R3 | 11C_JH vs 25C_EtOH | 2.78 | 1.63e−02 | * |
| Dunn (BH) | R3 | 11C_JH vs 25C_JH | 4.319 | 9.42e−05 | *** |
| Dunn (BH) | R3 | 25C_JH vs 25C_EtOH | −1.937 | 6.32e−02 | ns |
| Dunn (BH) | R4 | 11C_EtOH vs 25C_EtOH | −2.025 | 8.57e−02 | ns |
| Dunn (BH) | R4 | 11C_EtOH vs 25C_JH | 1.58 | 1.71e−01 | ns |
| Dunn (BH) | R4 | 11C_JH vs 11C_EtOH | −0.301 | 7.63e−01 | ns |
| Dunn (BH) | R4 | 11C_JH vs 25C_EtOH | −2.269 | 6.98e−02 | ns |
| Dunn (BH) | R4 | 11C_JH vs 25C_JH | 1.214 | 2.70e−01 | ns |
| Dunn (BH) | R4 | 25C_JH vs 25C_EtOH | −3.802 | 8.61e−04 | *** |
| Dunn (BH) | R5 | 11C_EtOH vs 25C_EtOH | 2.53 | 1.71e−02 | * |
| Dunn (BH) | R5 | 11C_EtOH vs 25C_JH | 5.144 | 1.62e−06 | *** |
| Dunn (BH) | R5 | 11C_JH vs 11C_EtOH | −2.95 | 6.45e−03 | ** |
| Dunn (BH) | R5 | 11C_JH vs 25C_EtOH | −0.726 | 4.68e−01 | ns |
| Dunn (BH) | R5 | 11C_JH vs 25C_JH | 1.93 | 6.43e−02 | ns |
| Dunn (BH) | R5 | 25C_JH vs 25C_EtOH | −2.945 | 6.45e−03 | ** |
| Wilcoxon rank−sum | R1 | 11C vs 25C | 25.5 | 3.57e−05 | *** |
| Wilcoxon rank−sum | R1 | EtOH vs JH | 157.5 | 8.68e−01 | ns |
| Wilcoxon rank−sum | R2 | 11C vs 25C | 68.5 | 4.51e−03 | ** |
| Wilcoxon rank−sum | R2 | EtOH vs JH | 44.5 | 2.51e−04 | *** |
| Wilcoxon rank−sum | R3 | 11C vs 25C | 55.5 | 1.13e−03 | ** |
| Wilcoxon rank−sum | R3 | EtOH vs JH | 160.5 | 1.00e+00 | ns |
| Wilcoxon rank−sum | R4 | 11C vs 25C | 171.5 | 4.84e−01 | ns |
| Wilcoxon rank−sum | R4 | EtOH vs JH | 59 | 2.19e−03 | ** |
| Wilcoxon rank−sum | R5 | 11C vs 25C | 57 | 1.42e−03 | ** |
| Wilcoxon rank−sum | R5 | EtOH vs JH | 28 | 8.70e−06 | *** |

### DI+ Mean Proportion

| Test | Subgroup | Comparison | Statistic | p (BH−adj) | Sig |
|---|---|---|---|---|---|
| Dunn (BH) | All regions | 11C_EtOH vs 25C_EtOH | 3.345 | 2.46e−03 | ** |
| Dunn (BH) | All regions | 11C_EtOH vs 25C_JH | 5.263 | 8.49e−07 | *** |
| Dunn (BH) | All regions | 11C_JH vs 11C_EtOH | −1.795 | 8.73e−02 | ns |
| Dunn (BH) | All regions | 11C_JH vs 25C_EtOH | 1.257 | 2.09e−01 | ns |
| Dunn (BH) | All regions | 11C_JH vs 25C_JH | 3.232 | 2.46e−03 | ** |
| Dunn (BH) | All regions | 25C_JH vs 25C_EtOH | −2.338 | 2.91e−02 | * |
| Kruskal−Wallis | All regions | All 4 groups | 29.301 | 1.94e−06 | *** |
| Wilcoxon rank−sum | All regions | 11C vs 25C | 16 | 3.28e−07 | *** |
| Wilcoxon rank−sum | All regions | EtOH vs JH | 73 | 4.87e−03 | ** |

### Ovary Status

| Test | Subgroup | Comparison | Statistic | p (BH−adj) | Sig |
|---|---|---|---|---|---|
| Fisher's exact (BH) | Diapaused | 11C_EtOH vs 25C_EtOH | − | 1.01e−12 | *** |
| Fisher's exact (BH) | Diapaused | 11C_EtOH vs 25C_JH | − | 1.34e−12 | *** |
| Fisher's exact (BH) | Diapaused | 11C_JH vs 11C_EtOH | − | 1.20e−06 | *** |
| Fisher's exact (BH) | Diapaused | 11C_JH vs 25C_EtOH | − | 6.22e−04 | *** |
| Fisher's exact (BH) | Diapaused | 11C_JH vs 25C_JH | − | 9.68e−04 | *** |
| Fisher's exact (BH) | Diapaused | 25C_JH vs 25C_EtOH | − | 1.00e+00 | ns |
| Logistic regression (BH) | Diapaused | (Intercept) | 3.445 | 1.14e−03 | ** |
| Logistic regression (BH) | Diapaused | treatment_cd25C | −0.009 | 9.99e−01 | ns |
| Logistic regression (BH) | Diapaused | treatment_cd25C:treatment_drugJH | 0.001 | 9.99e−01 | ns |
| Logistic regression (BH) | Diapaused | treatment_drugJH | −3.533 | 1.14e−03 | ** |

### LOS

| Test | Subgroup | Comparison | Statistic | p (BH−adj) | Sig |
|---|---|---|---|---|---|
| Fisher's exact (BH) | LOS | 11C_EtOH vs 25C_EtOH | − | 7.77e−04 | *** |
| Fisher's exact (BH) | LOS | 11C_EtOH vs 25C_JH | − | 4.17e−02 | * |
| Fisher's exact (BH) | LOS | 11C_JH vs 11C_EtOH | − | 1.80e−01 | ns |
| Fisher's exact (BH) | LOS | 11C_JH vs 25C_EtOH | − | 1.18e−05 | *** |
| Fisher's exact (BH) | LOS | 11C_JH vs 25C_JH | − | 7.77e−04 | *** |
| Fisher's exact (BH) | LOS | 25C_JH vs 25C_EtOH | − | 2.30e−01 | ns |
| Logistic regression (BH) | LOS | (Intercept) | −0.684 | 9.87e−01 | ns |
| Logistic regression (BH) | LOS | treatment_cd25C | −0.012 | 9.92e−01 | ns |
| Logistic regression (BH) | LOS | treatment_cd25C:treatment_drugJH | 0.011 | 9.92e−01 | ns |
| Logistic regression (BH) | LOS | treatment_drugJH | 1.551 | 4.84e−01 | ns |
| Logistic regression 11C only (BH) | LOS | (Intercept) | −0.684 | 4.94e−01 | ns |
| Logistic regression 11C only (BH) | LOS | treatment_drugJH | 1.551 | 2.42e−01 | ns |
